# A stable pollination environment limits current but not potential evolution of floral traits

**DOI:** 10.1101/581827

**Authors:** Maria Clara Castellanos, Javier Montero-Pau, Peio Ziarsolo, Jose Miguel Blanca, Joaquin Cañizares, Juli G. Pausas

## Abstract

The vast variation in floral traits at a macroevolutionary level is often interpreted as the result of adaptation to pollinators. However, studies in wild populations often find no evidence of pollinator-mediated selection on flowers. Evolutionary theory predicts this could be the outcome of long periods of stasis under stable conditions, followed by shorter periods of pollinator change that provide selection for innovative phenotypes. We asked if periods of stasis are caused by stabilizing selection, absence of other forms of selection on floral traits, or by low trait ability to respond even if selection is present. We studied *Ulex parviflorus*, a plant predominantly pollinated by one bee species across its range. We measured heritability and evolvability of floral traits, using genome-wide molecular relatedness in a large wild population, and combined this with estimates of selection on the same individuals. We found evidence for both stabilizing selection and low trait heritability as explanations for stasis in flowers. The area of the standard petal is under stabilizing selection, but the variability observed in the wild is not heritable. A separate trait, floral weight, in turn presents high heritability, but is not currently under selection. We show how a stable environment can lead to a lack of evolutionary change, yet maintain heritable variation to respond to future selection pressures.

## Introduction

Flowering plants exhibit a striking diversity in floral form and function, and because flowers are reproductive organs, the causes and dynamics of their evolution are crucial for understanding plant biodiversity. Much of the variation in floral traits at a macroevolutionary level is often interpreted as the result of adaptations to pollinators (Fenster et al. 2004). Experimental studies also confirm that many floral traits can be subject to selection by pollinators (reviewed by Parachnowitsch and Kessler 2010, Caruso et al. 2018). However, field studies measuring pollinator-mediated evolution of floral traits often find erratic evidence for strong selection in wild populations (Harder and Johnson 2009). In their review, Harder and Johnson found that only about 1/3 of the studies reported significant selection on floral traits. A possible reason for this paradox is the prevalence of periods of stasis, where pollinator-mediated selection on flowers is relaxed or limited to stabilising selection under stable conditions, interrupted by more unstable periods where pollinator changes can provide selection for innovative phenotypes (e.g. Galen 1989, Harder and Johnson 2009, Mackin et al. 2021). The causes of evolutionary stasis, in this case for flowers but common in many types of traits and organisms, is one of the long-standing questions around stasis that have intrigued evolutionary biologists (Merilä et al. 2001, Estes and Arnold 2007).

Evolutionary theory’s most basic prediction in quantitative trait evolution is captured by the breeder’s equation (R = *S* * *h^2^*) where, for an evolutionary response to take place (R), phenotypic traits need to be under directional selection (*S*) and harbour enough heritable variation for evolution to take place in the wild (*h^2^*; Falconer and Mackay 1996). The multivariate extension of the breeder’s equation reflects the fact that traits do not evolve in isolation and predicts a response determined by selection on multiple traits and a matrix of genetic variance (Lande and Arnold 1983). In any of its forms, the breeder’s equation has been found to be too simplistic to consistently explain periods of stasis (“the missing response to selection”) in many types of traits in wild animal and plant populations, suggesting that several other mechanisms are also involved (see Merilä et al. 2001 and Pujol et al. 2018). However, our knowledge on even the fundamental aspects of the nature of selection and the presence of heritable variation in wild populations is still limited, particularly for plants. The breeder’s equation is thus still useful by providing a good starting set of predictions, where periods of stasis can be the consequence of stabilizing or a lack of directional/disruptive selection on traits, or they can also be the result of low levels of heritable variation even if selection is present.

In the case of flowers, an appropriate model to study the role of these two non-exclusive scenarios is a plant with a stable single dominant pollinator. Under these stable conditions, floral traits can be expected to experience low levels of pollinator-driven selection, even if heritable variation is maintained in the population. Heritable variation in floral traits has been shown for many species in greenhouse studies (reviewed in Ashman and Majetic 2006, Opedal 2019), as well as in a few field studies (Schwaegerle and Levin 1990, Mazer and Schick 1991, Campbell 1996, Galen 1996). If heritable variation is generally present, stabilizing selection, or a relaxation of selection of any kind could be the most likely explanation for stasis in floral traits in populations with stable environments. Stabilizing selection is one the main mechanisms invoked to explain periods of stasis; however, it is often hard to detect in microevolutionary studies (Haller and Hendry 2013).

However, trait heritability in wild conditions could also be lower than the estimates under artificially reduced environmental variation. Traditional greenhouse and common garden studies of heritability allow for control of local environments and genetic background, but heritability values measured under controlled conditions can be systematically overestimated compared to wild conditions (Conner et al. 2003, Winn 2004). This can be caused by higher environmental variability in the field, as well as decreased expression of additive variance, or potential differences in survival in the field compared to the greenhouse, all leading to smaller heritability estimates. The alternative of measuring heritability directly in the field, although being more realistic, was until recently constrained by difficulties in either designing complex crossing and planting experiments (see Campbell 1996), or establishing relatedness among individual plants growing in the wild. This has now changed thanks to access to large and highly informative molecular markers that are distributed over the entire genome (Castellanos et al. 2011, Stanton-Geddes et al. 2013). Using genome-wide markers to measure genetic similarity of plants growing in the wild (in the form of a genome-wide relatedness matrix, *GRM*), makes it possible to estimate the proportion of the phenotypic covariance that is explained by relatedness (i.e. heritability) in the focal trait (Ritland 1996). This approach can incorporate environmental factors in the statistical estimation of heritability, to provide us with an ecologically realistic view of what plant populations are experiencing in natural conditions and help us understand the role of standing genetic variation in evolution (Campbell 1996, Kruuk et al. 2014).

We study the consequences of a stable pollination environment on natural selection and the heritability of floral traits by focusing on a plant with a single dominant pollinator, the Mediterranean gorse (*Ulex parviflorus*). *Ulex* and relatives (the large legume subfamily Faboidae) have complex irregular butterfly-type flowers (“papilionoid” or “keel” flowers; Fig. 1) believed to be specialized on bee pollination, with traits that both enhance pollinator attraction and mechanical interactions that improve pollination success (Westerkamp 1997). Over long periods of time, the evolution of flowers in this lineage was thus very likely shaped by adaptation to its pollinators. In contemporary timescales, observations across the current distribution of *Ulex parviflorus* show that honey bees (*Apis mellifera*) are currently the prevailing pollinator, with consistently low visitation rates, including in areas with low human influence. Dominance of honey bee visitation was observed by Herrera (1988; no precise percentage of visits reported) and again decades later by Reverté et al. (2016; 63% of visits were by honey bees) in coastal populations in southern and eastern Spain respectively, and has also been observed in inland populations in Cazorla, Spain (93% of visits; C.M. Herrera pers com.). In this currently stable system, we predict 1) a relaxation of directional or disruptive selection on floral traits and predominance of stabilizing selection if any type of selection is present, and/or 2) low trait heritability as a consequence of selection in the past that might have reduced genetic variation.

**Figure 1.**
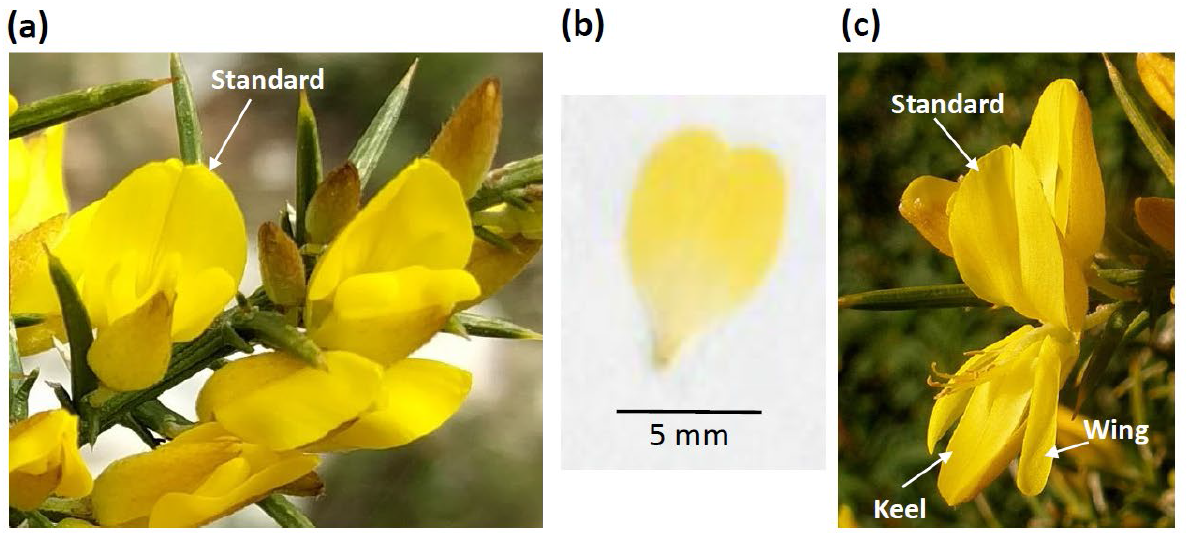
Flowers of *Ulex parviflorus*. (a) Flowers previous to a visit with standard petal extended and reproductive organs enclosed by the keel petals and calix. (b) Pressed standard petal. (c) Flower after being “triggered” by a bee visit, showing all petals and exposed reproductive organs. (Photo credits: (a) MC Castellanos, (c) J. Quiles).

To test these predictions, we measured trait heritability and natural selection on the same plant individuals in a wild population; this allowed us to assess the potential for evolution in response to current and future selection. To our knowledge, this is the first time this approach is used successfully to study floral traits in the wild. We measured floral morphology and pollinator visits, along with estimates of natural selection, genetic correlations, evolvability, and heritability of two floral traits to 1) determine if floral traits in a stable pollination environment are currently potentially evolving in response to selection, and 2) if not, to establish if the causes for a lack of evolutionary response are related to the nature of selection, low heritability, or both. With this we aim to contribute to understanding the underlying causes of the missing response to selection in natural plant populations.

## Materials and Methods

### Study species and sampling locations

*Ulex parviflorus* Pourr. (Mediterranean gorse; Fabaceae) is a thorny perennial shrub that lacks true leaves in the adult stage and grows up to 2 m. Species in the genus *Ulex* have yellow hermaphroditic flowers visited and pollinated by large-bodied bees, similar to other species in the tribe Genistae (Herrera 2001). Flowers do not produce nectar, and the bees visit to collect pollen, but to do so, they need to be heavy enough to actively trigger the explosive mechanism for pollen release. Reproductive organs are enclosed by specialized petals, the keel and the wings (Fig. 1). When an insect presses the keel with its hind legs, the previously concealed stamens and stigma are released with a powerful upward movement, placing a cloud of pollen grains on the ventral side of the bee. After a visit flowers do not recover their original shape, with stamens and style now protruding from the keel, and are rarely visited by large bees again, but can receive visits from smaller insects like hoverflies and solitary bees. *Ulex parviflorus* is self-compatible but depends on pollinators to set fruit (Herrera 1987). Flowering starts in the winter and can last for a few months into spring.

The species is widespread along the western Mediterranean coast from Southern France to southern Portugal. It is a successful colonizer of oldfields resulting from abandoned human activities, as well as recently burnt areas, thanks to numerous adaptations to recruitment after fire (Pausas et al. 2012, Pausas and Moreira 2012, Pausas et al. 2017). The seeds form a persistent bank in the soil, where they remain dormant until the heat produced during a fire breaks dormancy and stimulates germination in post-fire conditions (Moreira et al. 2010). Current landscapes in eastern Spain are a mosaic of oldfields and postfire shrubland (Pausas & Millán 2019), where *Ulex parviflorus* is very abundant and distributed continuously from the lowlands up to 900m of altitude (Fig S1 in supplementary materials). As a consequence of the high connectedness, there is very low genetic differentiation in the study area (Moreira et al. 2014; see also Supplementary methods and Fig. S3), and different stands cannot be considered distinct populations. For our sampling, we selected six sites within this continuous population, aiming to capture the potential variability of mature *U. parviflorus* stands in the area (Table S1, Fig. S1). By sampling at different altitudes, for example, we include variability in floral traits along the elevational gradient (Fig. S2 in supplementary materials). At each site, we tagged 40 individual plants (240 plants in total) for phenotypic and genotypic characterization as described below. Individuals were at least 5 m apart from each other and blooming at the time of sampling.

### Pollinator censuses

To quantify the diversity of floral visitors as well as visitation rates, we ran multiple three-minute pollinator censuses at different times of the day, for up to five hours of observations per locality, on two separate days during peak blooming in 2014 (plus extra censuses in two localities in 2013). Each census recorded the number and identity of visitors to patches of flowers on haphazardly chosen shrubs, including but not limited to the 40 tagged individuals. We counted the number of flowers surveyed in each census and the number of flowers visited to estimate the per-flower visitation rate.

### Floral phenotypes

We collected five haphazardly selected flowers from each individual plant for phenotypic characterization of two floral traits that function as proxies for flower showiness and flower size. The area of the upwards-facing petal, or standard, plays a key role in flower showiness, as it is the largest and most visible petal in these typical papilionoid flowers (Fig. 1; standard petals are often called flag or banner petals). We removed standards from all flowers when fresh, and pressed them flat individually in a plant press until dry. We then used scanned images of the standards to measure their surface area with the Image-J analysis software (Schneider et al. 2012).

Flower mass reflects the size of the flower and is important in the Genistae as it determines the size of the insects that can visit the flowers (Herrera 2001, Córdoba and Cocucci 2011). It was estimated as the dry weight of flowers (calyx and corolla) after removing the standard petal and the pedicel, and carefully brushing off all pollen grains. Flowers were pressed and oven-dried at 40°C for 48 hours and weighed to the nearest 0.01 mg.

These above traits were chosen because they can be expected to play an important role in the interaction with pollinators and thus be under pollinator-driven natural selection (see Study Species above). As is the case in many complex flowers, the two traits studied are likely to co-vary, and analyses below are designed to take this into consideration. We have no reason to suspect that there is variability in these traits with flower age (see Herrera 2001). We have never observed florivory in this species and thus assume that herbivores will not directly select for the two focal traits in this study.

### Plant genotyping

Fresh terminal twigs were collected from each tagged individual plant and dried in silica gel previous to DNA extraction. The extraction was performed using the Speedtools plant DNA extraction kit (Biotools, Madrid, Spain), with modifications to the manufacturer’s protocol to optimize DNA quantity and quality extracted for this highly lignified species. We used the Genotyping-by-Sequencing (GBS) protocol to identify single nucleotide polymorphisms (SNPs) across the genome (Elshire et al. 2011). Illumina libraries for our 240 individuals were constructed by digesting genomic DNA with a restriction enzyme. Two libraries were built for each plant after separate digestions with *PstI* and *EcoT22I*, in order to increase the number of high quality SNPs. Library construction and sequencing was performed by the Genomic Diversity facility at Cornell University (USA). SNP calling was implemented using the UNEAK pipeline (Lu et al. 2013) in the TASSEL v.3 software package (Bradbury et al. 2007), designed for data sets without a reference genome.

The final SNP dataset used for the analysis of relatedness below excluded loci that were genotyped in less than 90% of individual plants. The minimum allele frequency allowed to retain loci was set to MAF > 0.01. We also excluded individuals with low genotyping rates (under 85% of loci). After applying these filters, we further manually removed remaining loci with extreme values of observed heterozygosity (under 2% and higher than 98%), after estimating oHET with PLINK command “–Hardy” (Purcell et al. 2007).

### Fitness estimates and phenotypic selection

We estimated fruit set in the same 40 individual plants in each of the six localities as a proxy for female reproductive success. For this, we labelled a representative flowering twig in each plant during flowering peak. When fruits were already developing (browning capsules) a few weeks later, we collected the labelled twig in a paper envelope. Back in the laboratory we measured 10 cm of twig to calculate a) the number of fruits developing normally, and b) scars left by all flowers produced by the twig, clearly visible under a dissecting microscope. From this we calculated fruit set as the proportion of flowers that develop into a fruit. The majority of fruits had one (71% of 3200 fruits examined) or two seeds (25%), with a mean number of 1.22 seeds/fruit across all individuals.

We estimated selection parameters to test for both linear (directional selection) and non-linear (stabilizing or disruptive) selection on the two floral traits, using fruit set as the response fitness variable in the models. Because floral weight and standard area show a significant phenotypic correlation (even though floral weight did not include the standard, Pearson *r* = 0.43, P < 0.001), we estimated selection gradients in addition to selection differentials. Selection differentials provide univariate estimates of selection without considering other traits, while gradients provide estimates on correlated traits. By estimating the four selection parameters - standardized linear (S), and quadratic (c) selection differentials, and standardized linear (β) and quadratic (γ) selection gradients - we can explore direct and indirect selection on the floral traits.

We used generalised additive models (GAM) to measure selection parameters on absolute fitness values, following the approach developed by Morrisey and Sakreda (2013). This approach provides quantitative estimates of selection differentials and gradients for non-normal fitness components, testing for both linear and quadratic selection. We fitted GAMs for binomial fruit set data (fruits developed in relation to total flowers), using a logit link function and assuming a binomial error distribution with the *mgcv* package in R. We used univariate GAMs to estimate selection differentials, and included both floral traits into a bivariate model to estimate selection gradients. To control for local effects, we included locality as a random factor in all models. Models included additive spline effects on all factors. Differential and gradient parameters were estimated based on numerical approximations of first and second partial derivatives of relative fitness, averaged over the distribution of observed phenotype. To calculate the significance of selection differentials and gradients, we used the bootstrap approach (n= 1000 samples) implemented in the *gsg* package in R (Morrissey and Sakrejda 2013).

### SNP-based relatedness and quantitative genetic parameters

Pairwise relatedness between all pairs of individuals was estimated from the similarity of their SNP genotypes. To estimate GRM, the genome-wide relatedness matrix among all pairs of individuals, we used the realized relatedness method of VanRaden (2008) and Astle and Balding (2009) as implemented in the *kin* function of package *synbreed* in *R* (Wimmer et al. 2012; see details in Supplementary methods). Relatedness values under this approach are a measure of excess allele sharing compared to unrelated individuals. As a consequence, negative values can be common and correspond to individuals sharing fewer alleles than expected given the sample.

To estimate additive genetic variance (and then heritability and evolvability) we used a linear mixed ‘animal model’ approach to model the phenotypic variance in floral traits while including the variance explained by relatedness (Wilson et al. 2010). We included the elevation above sea level as a fixed effect to account for environmental variability among plants, because elevation is the main factor that varies among localities (Fig. S2) and this could affect floral traits as in other species (Herrera 2005). In addition to the additive genetic effects (see model below), models included two more random effects: the site of origin of each plant, to account for unmeasured local environmental effects that could co-vary with genetic variation, and the individual identity to account for intra-individual effects (a “permanent environment” effect in Wilson et al. 2010), because we had five flower replicates per plant. We ran a univariate model for each of the two floral traits studied, specified as:

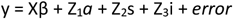

where y is the vector of floral trait values, β is the vector of fixed effects (with X as the incidence matrix), Z_1_, Z_2_ and Z_3_ are incidence matrices for the random effects *a* (individual identity to partition additive genetic effects), s (the locality), I (individual identity to model intra-individual effects caused by differences among replicate flowers from the same individual), and e is the residual error. The variance-covariance structure of random factor *a* in the model is defined by GRM·V_*a*_, where GRM is the genome-wide relatedness matrix between plant pairs, and V_*a*_ is the additive variance to be estimated. To test for the effect of not including the spatial and environmental predictors in the models, we also ran a ‘naïve’ version of each model that included only the relatedness and individual effects (Castellanos et al. 2015). We ran Bayesian animal models using package *MCMCglmm* for *R* (Hadfield 2010) with both floral weight and standard petal area modelled as continuous traits. For modelling the standard area, we used parameter-expanded priors for the distribution of variance components following the *χ*^2^ distribution with one degree of freedom. Each analysis was iterated long enough to obtain 5000 independent chains (see supplementary methods and Table S2 for model details, scripts and prior selection).

Narrow sense heritability (*h^2^*) was then estimated as the proportion of the total phenotypic variance assigned to the individual (i.e. to the additive genetic variance, V_*a*_):

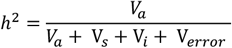

where V_S_ is the variance explained by the site of origin, V_*i*_ is the intra-individual variance in the trait, and V_error_ is the residual variance. We also estimated the narrow sense evolvability e, i.e. the mean-standardized additive genetic variance, *e* = V_*a*_ / *x*^2^, where *x* is the trait mean. *e* reflects the expected percentage of change of a trait under a unit strength of selection per generation (Houle 1992, Hansen et al. 2003) and provides an estimate of evolvability that is independent of trait variation and comparable across traits.

In addition, we estimated the genetic correlation (r_G_) between floral weight and standard area by running a bivariate animal model in *MCMCglmm*. In this case we used the same fixed and random factors as in the univariate models above (see supplementary methods for prior information).

Finally, we calculated the statistical power of our heritability estimates using the GCTA-GREML Power Calculator (https://shiny.cnsgenomics.com/gctaPower; Visscher et al. 2014). This analysis uses the GCTA approach by Yang et al. (2011), also a population-based estimation method of heritability that differs from our models in using unrelated individuals only. Even so, it can provide an indirect power estimation for this study.

## Results

### Pollinators

We recorded 364 visits to 22522 censused flowers in 28 hours of observations across the six *U. parviflorus* localities. Of those, 331 (92%) were visits by the honeybee *Apis mellifera*. Further 25 visits were by *Bombus sp*. individuals (7%). The remaining 3 visits were to already open flowers by small Coleoptera and a hoverfly, both unlikely to contact stigmas and carry out pollination. Across sites, we found an average visitation rate of 0.015 (±0.057) visits per 3-minute census to an individual flower, which translates into one visit every 3.3 hours, on average. Visitation rates were similar across localities, except for one where visits were significantly more frequent (Simat average visitation rate= 0.03 visits per census).

### Floral phenotypes and selection

Flowers showed variation in the two traits measured, flower weight and standard petal area, both within and across localities. A variance partition analysis showed that the variance in both traits across the five flowers sampled per plant was small (under 15% of the total variance), so we ran the selection analysis below using mean floral values for each individual plant (see also Herrera 2001). The coefficient of variation (CV) of these mean values in our data set and found that they show similar values (flower weight CV =21.1%, standard petal area CV = 21.2%).

We found no evidence of linear directional selection on floral traits, either in univariate models (s coefficients) or models of correlated selection incorporating both floral variables (β coefficients, Table 1). However, we found evidence for univariate quadratic effects in both traits (c coefficients), and quadratic gradients (γ coefficients) for standard area. This suggests that floral weight is not under direct selection, while there is strong evidence for stabilising selection on standard petal area (Fig. 2).

**Table 1.** Directional and quadratic selection coefficients (± standard errors) for the two floral traits studied.

| Trait | Differential |  | Gradient |  |
| --- | --- | --- | --- | --- |
| | Directional, S | Quadratic, c | Directional, $\beta$ | Quadratic, $\gamma$ |
| Standard petal area | -0.003 ± 0.003 ns | **-0.002 ± 0.000 \*\*\*** | -0.041 ± 0.039 ns | **-0.102 ± 0.029 \*\*\*** |
| Flower weight | -0.018 ± 0.033 ns | **-0.070 ± 0.023 \*\*** | -0.022 ± 0.047 ns | -0.038 ± 0.021 ns |
| Interaction | | | | 0.000 ± 0.002 ns |

**Figure 2.**
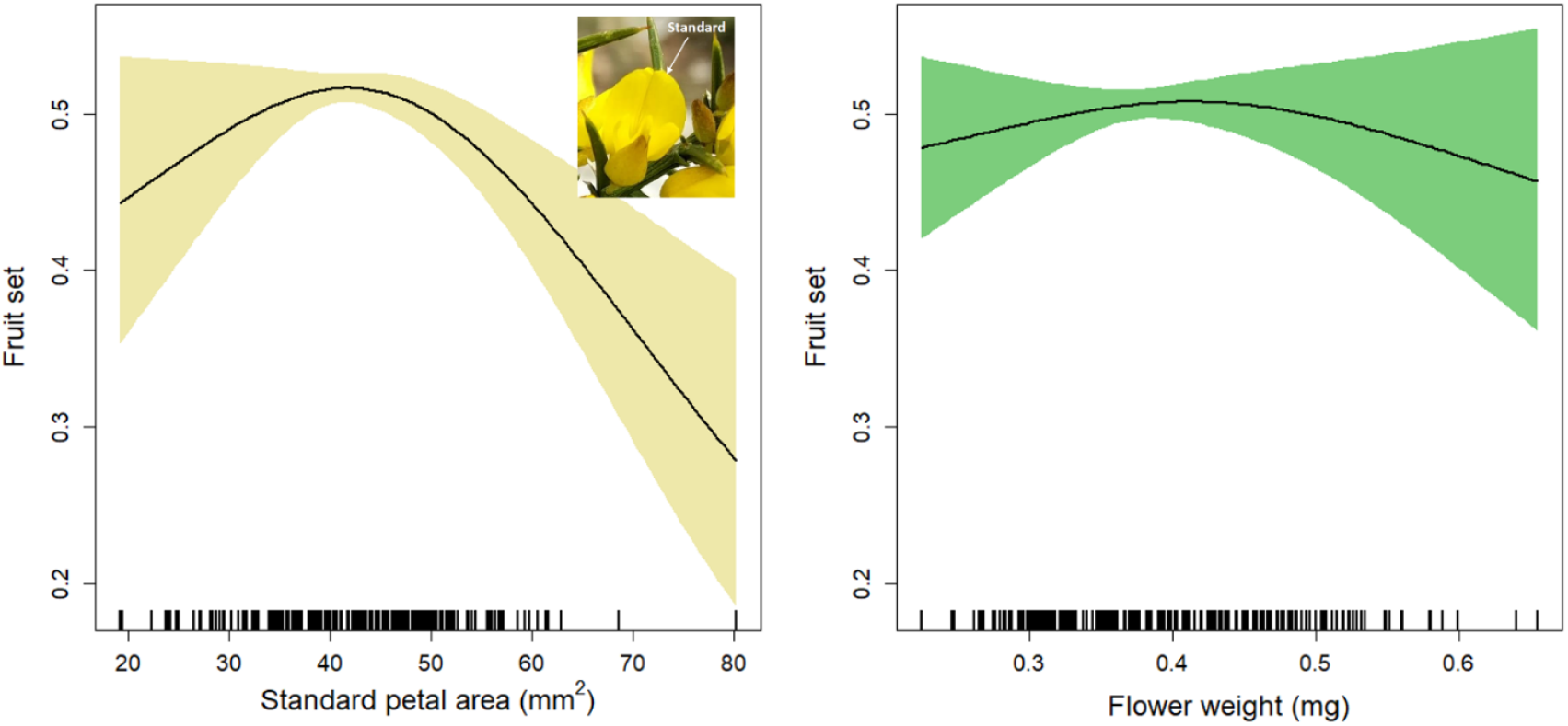
Fruit set as a function of the two floral traits measured, (a) standard petal area and (b) flower weight (a proxy for floral size). Lines are univariate generalised additive model fits with shaded areas showing 95% confidence interval

### Genomic markers and population genetic structure

The GBS sequencing approach yielded a large number of polymorphic SNPs across individuals (261,775 SNPs before quality filtering). After MAF and heterozygosity filtering, we retained 10,421 high-quality SNPs that were present in at least 90% of individuals across all localities. The analyses below use this dataset to estimate genomic relatedness; however, we also tested for the effect of retaining a larger number of SNPs (with presence in at least 50% of the individuals, which leads to a higher number of genotypes imputed by *synbreed*, see Supplementary methods). Analysis with this larger dataset produced the same qualitative results, so that retaining more (but highly imputed) markers did not add valuable information on the relatedness among our study plants. Therefore, all analyses below use the smaller dataset with 10,421 SNPs.

### Heritability, evolvability and genetic correlation

Pairwise relatedness among sampled individuals varied markedly and was overall relatively low (average values ranging from −0.09 to 0.79, but with most values <0.2), even within localities (Fig. S4); this supports the prevalence of outcrossing in this species. The low population genetic structure and the presence of variance in relatedness provide the conditions for a reliable estimation of heritability in the field (Ritland 1996).

We found significant estimates of heritability and evolvability in flower weight (*h^2^* = 0.14, *e* = 0.42%; Table 2). For standard area, our models instead detected very low additive variance, yielding very low *h^2^* and *e* in this case (*h^2^* = 0.001, *e* < 0.001%; Table 2). For both traits, Deviance Information Criterion (DIC) values for the heritability naïve models were larger than for the complete model (Table S2), indicating a better fit for the latter. The naïve models included only the relatedness among individuals and neither environmental nor spatial predictors, and showed estimated *h^2^* values substantially higher than our final estimates (Table 2).

**Table 2.**
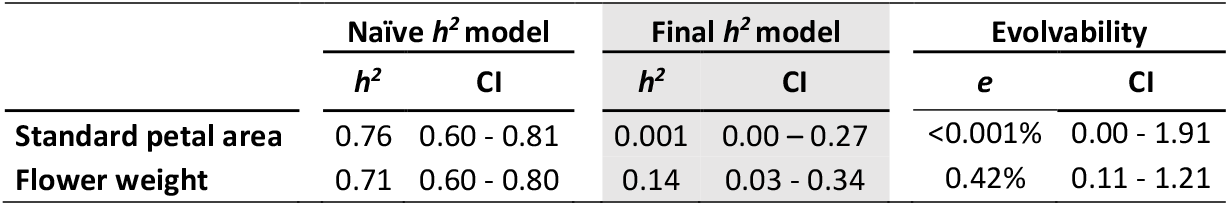
Estimates of heritability *h^2^* and evolvability *e* (with 95% credibility intervals, CI) for floral traits in wild *Ulex parviflorus*. ‘Naïve’ heritability models did not include spatial or environmental predictors.

| | Naïve $h^2$ model | | Final $h^2$ model | | Evolvability | |
| --- | --- | --- | --- | --- | --- | --- |
| | $h^2$ | CI | $h^2$ | CI | $e$ | CI |
| **Standard petal area** | 0.76 | 0.60 - 0.81 | 0.001 | 0.00 - 0.27 | <0.001% | 0.00 - 1.91 |
| **Flower weight** | 0.71 | 0.60 - 0.80 | 0.14 | 0.03 - 0.34 | 0.42% | 0.11 - 1.21 |

Our sample size for estimating the variance components are limited (240 plants) and thus the statistical power was low to detect very low values of heritability (power = 0. 09 for *h^2^* values of 0.1; see full analysis in Supplementary Table S3). Even with this limited power, we were able to estimate a significant *h^2^* value for flower weight in this population. Our bivariate analysis found a low genetic correlation between the two floral traits that is indistinguishable from zero (r_G_ = 0.06). Credible intervals were large (−0.139 to 0.381), so we cannot support the presence of a genetic correlation.

## Discussion

We provide an example of a stable environment that has led to stasis in phenotypic traits, yet maintaining enough heritable variation for responding to possible novel selection pressures, at least in some traits. This follows general theoretical expectations and exemplifies two of the main expected biological explanations for stasis (Merilä et al. 2001), with evidence for both stabilizing selection and low trait heritability as alternative explanations for the potential lack of evolution. Our study focuses on flower traits, but the findings can be relevant to any trait under stable conditions, in the context of the missing response to selection. In *Ulex parviflorus*, the area of the standard petal is currently under stabilizing selection, but the variability we observe in the field is not heritable. Floral weight, in turn, presents high heritability, but is not currently under selection.

Stable pollinator communities are potentially a common feature of many plant species under stable environmental conditions. A low diversity in pollinators does not necessarily imply stability in terms of selection, because few pollinator species that are functionally different (e.g. belong to different taxonomic groups) and vary in space or time, could provide increased opportunities for selection. In the case of *Ulex parviflorus*, however, current evidence shows that honey bees are the dominant pollinators in all surveyed populations, including the one studied here and other distant localities, often sampled decades apart (Herrera 1988, Reverté et al. 2016, C.M. Herrera pers com). Other species in the genus, including *U. europaeus, U. minor* and *U. galli*, are visited by a higher diversity of large bees (several species of *Bombus* and *Andrena*; Kirchner and Bullock 1999, Bowman et al. 2008, Falk 2011). The dominance of honey bees in *U. parviflorus* populations could be seen as a consequence of the large anthropogenic influence across its range; however, populations in an area with low human influence and high pollinator diversity (Sierra de Cazorla, see Herrera 2019) corroborate the dominance of honey bees as pollinators of this species across its current distribution in contemporary time scales. Regardless of the reasons for low pollinator diversity, all this suggests that the opportunity for selection is overall stable for *U. parviflorus* and our study provides evidence on how this stability can lead to lack of current evolution in floral traits. On the opposite side of the spectrum, field studies that do detect directional selection on unmanipulated floral traits often focus on plants that are exposed to changing pollinator environments, either in different parts of the species range (Herrera et al. 2006, Anderson et al. 2010, Mackin et al. 2021) or in hybrid contact zones where there is selection against hybrid phenotypes (Campbell et al. 2018). Taken together, current evidence supports the idea that pollination-driven floral evolution takes place mostly during evolutionarily innovative periods driven by changing pollinators.

Stabilizing selection is one of the main potential causes of stasis in wild populations, and is also expected to be important in the particular case of floral traits that influence the accuracy of pollen deposition during the flower-pollinator interaction (Cresswell 2000, Armbruster et al. 2009). It is difficult to assess how common stabilizing selection is on floral (or many other) traits in wild plant populations, because studies rarely measure non-linear selection (Harder and Johnson 2009, Caruso et al. 2018), in addition to the overall difficulty of detecting it (Haller and Hendry 2013). For the standard petal in *Ulex*, we detected stabilizing selection for intermediate surface area. The size of this “flag” petal is expected to play an important role in pollinator attraction by increasing the floral colour display (Fig. 1), so that selection against smaller sizes can be expected. Too large standard petals could be selected against if they incur a higher cost for the plant. This cost could be even higher if large standard petals are developmentally restricted to overall larger flowers; however, our preliminary genetic correlation estimates suggest only a weak association between standard petal area and floral weight. This is consistent with a previous study that carefully dissected the role of the different petals in another keel flower; in *Collaea argentina*, Córdoba et al. (2015) found that the standard petal is not functionally integrated with another set of floral traits that collectively regulate the enclosing mechanism of stamens and pistil. That is, the mechanics of protecting the enclosed rewards in these flowers can be independent of pollinator attraction as we expected, and selection can vary across floral parts.

Floral morphological traits often present high levels of additive genetic variation (reviewed by Ashman and Majetic 2006, Opedal 2019); however, most of the studies in these reviews were performed in controlled environments where *h^2^* estimates are not directly relevant to evolution in the wild. Our field estimates of heritability fall within the lower range of those summarized in Figure 1 of Altman and Majetic (2006), as expected for field estimates, where environmental variation is higher. We found that flower weight shows significant heritability, but there was no detectable heritability in the standard petal area, that is in turn under stabilising selection. In the case of the latter, we cannot completely rule out that heritability is present but very small, because our sample size of 240 plants provides low statistical power to detect very small values of *h^2^*. Comparing petals in papilionoid flowers, Herrera (2001) found that the standard had higher phenotypic variance than other petals across Genisteae, and argued that its role in pollination was smaller than for the keel petals, in a similar way as Córdoba et al. (2015). This and our results suggest that this petal might be prone to high environmentally induced variation. It is also likely that long-term stabilizing selection has reduced the additive genetic variance in this trait, leading to the low *h^2^* values we detected, and consistent with the theoretical expectations for variance reductions under stabilising selection (although this is not always the case; Johson and Barton, 2005).

Heritability estimates have been criticised as poor standardized measures of evolutionary potential in realistic ecological settings, in part because of the covariance between environmental and genetic effects (Houle 1992, Hansen et al. 2011). In this study, we estimate heritability directly in the field, statistically controlling for environmental variation, and in the same individuals as those used to estimate natural selection. In this context, field heritability estimates provide a useful approach to understand the current evolutionary potential at the population level, because we are interested in the role of environmental effects on the phenotypic variance, as exposed to natural selection. An alternative measure of evolutionary potential, evolvability, uses the mean of trait values to standardize the additive genetic variance (as opposed to standardizing by the total phenotypic variance) and provides a comparable estimate of proportional change in a trait value after selection (Hansen et al. 2003). Our estimates of evolvability are within the range of values estimated for floral size across plant species (Opedal 2019). These values confirm our findings for heritability, that is, near-zero evolutionary potential for the standard petal area, but higher values for flower weight. In the latter case, evolvability is estimated to be significant but small (under 1% of the trait mean value), suggesting that change in this trait would be slow unless submitted to strong selection.

Our estimate of genetic correlation between the two focal traits needs to be interpreted with caution as our sample size was relatively low. However, the lack of a genetic correlation is not surprising given that we cannot detect significant additive genetic variation in one of the traits (the area of the standard petal). This contrasts with the fact that there is a significant phenotypic correlation between the two traits (although relatively low r = 0.43), but as suggested by previous studies, phenotypic correlations are not always good predictors of genetic correlations, even in highly integrated organs as flowers (Gómez et al. 2009). Again, this is consistent with the decoupling of petals found in a related species with keel flowers (Córdoba et al. 2015). It is thus possible that the phenotypic correlation is caused by shared environmental factors that affect both traits in *Ulex* flowers, further emphasizing the importance of studying evolutionary potential in field conditions.

Even though we could not detect a genetic correlation between the two floral traits studied here, our analysis does not include selection on other (unmeasured) potentially correlated traits. Another potential caveat is that our estimates of selection are based on fruit set alone, and we cannot rule out that the two focal traits might be under selection through the male function (van Kleunen and Burczyk 2008). However, the two traits studied here can be expected to affect pollen dispersal in similar ways as pollen deposition (and thus seeds sired), because the trigger mechanism forces both male and female reproductive organs to make contact with the bees at the same time. This means that factors affecting seed set and seed sire are probably highly related in keel flowers.

This study adds to a series of recent work using large sets of molecular markers to study quantitative genetics in wild populations, mostly focused on animals (Perrier et al. 2018), but also on plants (Castellanos et al. 2015). Studies comparing the accuracy of SNP-based relatedness matrices compared to pedigrees are consistently showing that they can be very good approximations, depending on the specie’s life history, and that they can provide higher analytical flexibility (Bérénos et al. 2014, Gienapp et al. 2017, Perrier et al. 2018). This is therefore an exciting time for studying the evolution of traits directly in the wild because field-based estimates of evolutionary potential provide new avenues to understand basic evolutionary questions (such as stasis and the role of plasticity in trait variation), but also the potential for wild organisms to respond to new selection pressures including those imposed by anthropogenic environmental change. In the case of flowers, the broad implications of our findings are that low-diversity pollination environments as those caused by anthropogenic pollinator declines can lead to reduced selection pressures, reduced opportunity for selection, and stasis, while exposure to new pressures could lead to novel evolutionary change.

## Final remarks

Our approach in this study attempts to capture the complex business of evolution in the wild assuming that measures of selection and heritability are sufficient and constant enough to predict microevolutionary change. This might not always be case, as other factors, such as a low a correlation between traits and fitness, genetic constraints, non-genetic inheritance, and plasticity, are not included in the breeder’s equation (reviewed in Pujol et al 2018). Nevertheless, in this study we provide a first set of results for a study system that are consistent with both theoretical predictions and observations in the field. Selection on floral traits is not restricted to pollinators alone (reviewed by Caruso et al. 2018), but regardless of the source of selection, our findings contribute to explain the macroevolutionary patterns of floral evolution where novel phenotypes are ubiquitous (exceptions are often related to very generalised pollination that is stable over evolutionary time, see Vasconcelos et al. 2019). Populations can experience periods of stasis, but heritable phenotypic variance can remain present in some traits. In combination with potential genetic correlations, this provides the potential to respond to novel selection. However, to fully understand evolutionary responses to rapid environmental change more studies in the wild are urgent.

