## Supporting Information for "A stable pollination environment limits current but not potential evolution of floral traits"

### Supplementary methods

#### Estimation of genomic relatedness

Previous to relatedness estimation, missing genotypic data were imputed using the *codeGeno* command in the *synbreed* package in R (Wimmer et al. 2012). This imputation step increased the number of available markers for analysis. Only slightly more than 1% of the genotypes (26,773 out of 2,344,725) were imputed.

Genomic relatedness was estimated with *synbreed* function *kin*, using the "realized" method. Here, GRM = ZDZ' where D is diagonal and given by Dii = 1/m[2∑pi(1-pi)]. This method weights markers by the reciprocals of their expected variance, and re-scales estimates of relatedness considering the current sample as the base population. The resulting GRM matrix is, by definition, positive semi-definite. To make matrices suitable for animal models, GRM matrices were forced to be positive-definite with the command *make.positive.definite* in package *corpcor* (Schäfer et al. 2014). This command computes the nearest positive definite symmetric matrix, using the algorithm by Higham (1988). This transformation adds very small values to the original relatedness estimates, e.g. the addition of a values around $3*10^{-13}$, and has a negligible effect on the matrix structure.

#### Estimation of variance components in *MCMCglmm* in R (Hadfield 2010)

The unweighted, inverted positive-definite matrix was then used to provide the covariance structure for the random 'animal' predictor in animal models for floral traits (input as *ginverse* in *MCMCglmm*). The GRM matrix is unweighted, as we have no preliminary information to assume that sampled SNPs are unequally linked to the genetic causal variants of the studied traits. We modelled floral weight and standard area as continuous traits using the default "gaussian" family distribution (see Table S2 for models and the main text for an explanation). Models for floral weight used the default inverse gamma distribution prior for the random effects, specified for three random effects as:

floral.weight.prior<- list (R = list (V = 1, nu = 0.002), G = list (G1 = list (V = 1, nu = 0.002), G2 = list (V = 1, nu = 0.002), G3 = list (V = 1, nu = 0.002)))

For the standard petal area, we obtained better posterior distributions using a parameter expanded prior with the $\chi^2$ distribution with 1 degree of freedom (see Gelman 2006):

standard.petal.prior<- list (R = list (V = 1, n = 0.002), G = list (G1 = list(V = 1, nu = 1, alpha.mu = 0, alpha.V = 1000), G2 = list (V = 1, nu =1, alpha.mu = 0, alpha.V = 1000), G3 = list (V = 1, nu =1, alpha.mu =0, alpha.V = 1000)))

To prevent any autocorrelation for variance components in MCMC chains, we ran MCMCglmm for 5 million iterations with a high thinning interval of 1000 after a burn-in of 100K chains. This resulted in an effective sample size of 5000 chains. Posterior modes and means of $h^2$ estimations were very similar to each other, as expected, and modes are reported throughout the paper.

For a bivariate model used to estimate genetic correlation we used the bivariate version of a non-informative inverse gamma distribution prior:

correlation.prior<-list(R=list(V=diag(2),n=1.002), G=list(G1=list(V=diag(2),n=1.002), G2=list(V=diag(2),n=1.002), G3=list(VV=diag(2),n=1.002)))

In this case, we ran MCMCglmm for 2 million iterations with a high thinning interval of 1000 after a burn-in of 100K chains, for a final sample size of 200 chains. The model was specified as the full models in Table S2 below, but including both floral traits.

**Within population genetic structure**

We tested for a potential regional genetic structure among localities in our sampled population by running the Bayesian clustering approach implemented in the software STRUCTURE v. 2.3.4 (Pritchard et al. 2000). This method assigns individuals to the optimal number of K genetic clusters based on allele frequencies at each locus. We truncated the dataset to 5K SNPs to keep running time manageable. We ran simulations including locality identifiers and with the LOCPRIOR option, to make sure that even weak genetic structuring could be detected. Simulation runs calculated the likelihood of clustering in K = 1 to 7 localities (one more than the actual sampled stands), and were run for $1 \times 10^4$ iterations after a $1 \times 10^5$ burn-in period, with separate values for alpha for each locality (alternative ancestry prior) as suggested by Wang (2017). Five runs were carried out for each value of K. The best value of K and summary figures were generated using the Clumpak server (Kopelman et al. 2015).

The analysis found weak population genetic structure among the study stands even when using our large set of molecular makers (5K SNPs), as expected for a widespread, effective colonizer species. A delta K analysis implemented in Structure Harvester (Earl and vonHoldt 2012) indicates an optimal number of populations around K=2, i.e., the minimum for the method (as it cannot favor K=1; Fig. S2). This confirms that gene flow is widespread (as was also concluded by Moreira et al. 2014 using AFLP markers) and that *Ulex* localities belong to a large population across the region.

**Supplementary Table and Figures**

**Table S1.** General characteristics of the six sampling sites (sorted by latitude), including name of municipality, geographical coordinates, elevation (in meters above sea level), mean annual temperature, and annual precipitation. The distance between sites ranged from 12 to 154 Km.

| Site | Latitude | Longitude | Elevation (masl) | T (ºC) | Precipitation (mm) |
|---|---|---|---|---|---|
| Ares del Maestrat | 40.41 | -0.08 | 820 | 14.4 | 760 |
| Sot de Chera | 39.60 | -0.92 | 775 | 14.2 | 600 |
| Chiva | 39.53 | -0.80 | 800 | 15.0 | 553 |
| Cheste | 39.52 | -0.62 | 170 | 17.7 | 422 |
| Montserrat | 39.39 | -0.58 | 190 | 16.7 | 490 |
| Simat de la Valldigna | 39.04 | -0.34 | 349 | 15.72 | 539 |

**Table S2.** Models and heritability estimates ($h^2$) for flower weight and standard petal area in *Ulex parviflus*, along with credible intervals (CI) and deviance information criterion values (DIC) for complete final models and "naïve" models with no spatial predictors (as in Table 3 in main text).

| Trait | model | | $h^2$ | CI | DIC |
|---|---|---|---|---|---|
| **Standard petal area** | full | spa ~ elevation, random= ind + site +ind2 | 0.001 | 0.00 – 0.27 | 5898.72 |
| | naïve | spa ~ 1, random= ind + ind2 | 0.76 | 0.60 - 0.81 | 5899.98 |
| **Flower weight** | full | weight ~ elevation, random= ind + site +ind2 | 0.14 | 0.03 - 0.34 | -4167.74 |
| | naïve | weight ~ 1, random= ind + ind2 | 0.71 | 0.60 - 0.80 | -4165.19 |

weight= floral weight; spa=standard petal area; ind= individual identifier (='animal') with covariance structure provided by their pairwise relatedness matrix, ind2= individual identifier used to account for the variance in floral traits within plants (n= 5 flowers per individual).

**Table S3.** Statistical power (probability of detecting heritability values >0) given our sample size of N=240 plants and an observed variance in relatedness of 0.00124, estimated from the variance of the diagonal of the relatedness GRM matrix. Power was estimated using the GCTA-GREML Power Calculator (https://shiny.cnsgenomics.com/gctaPower) with an error type 1 alpha= 0.5.

| $h^2$ | Power |
|---|---|
| 0.05 | 0.060 |
| 0.1 | 0.092 |
| 0.2 | 0.223 |
| 0.3 | 0.434 |
| 0.4 | 0.666 |
| 0.5 | 0.848 |

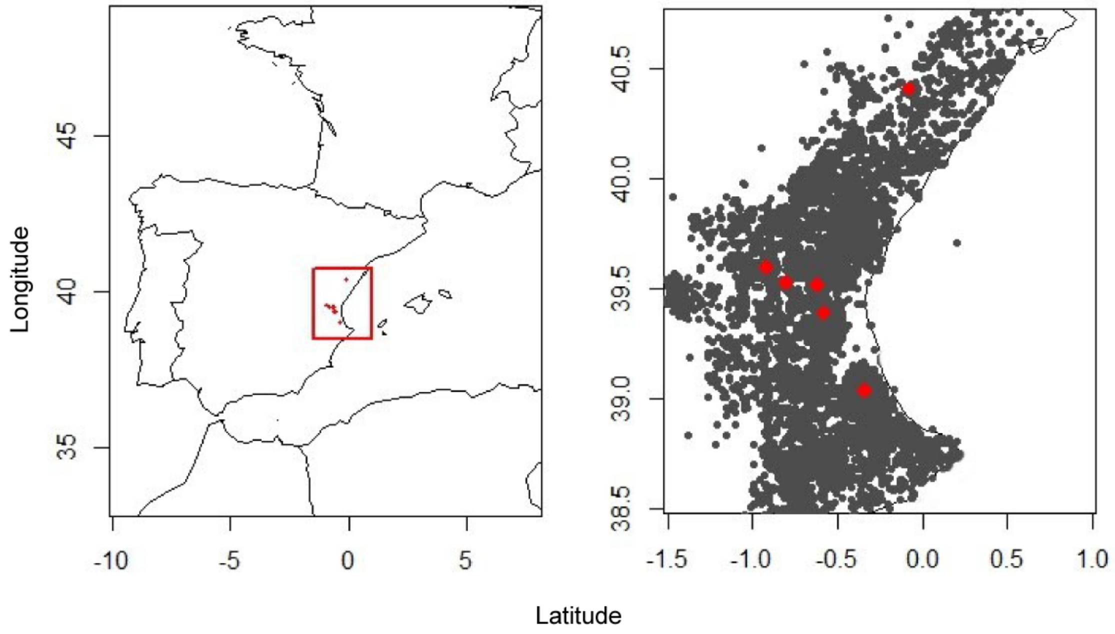


**Fig S1.** Distribution of *Ulex parviflorus* in the study region in Spain. The panel on the right corresponds to the region in the red square in the left panel. Dark grey dots are locations for all records for the species in GBIF.org (accessed 27 May 2019; doi: 10.15468/dl.h3sgmm); the actual distribution is likely to be even more widespread. Sampling sites are marked by the red dots.

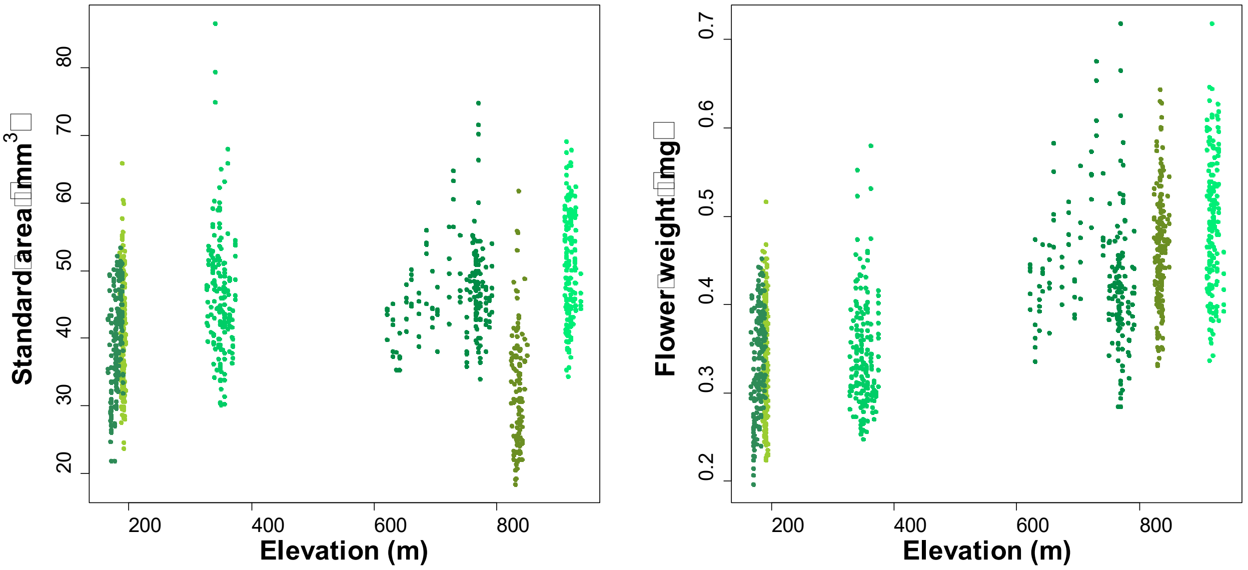


**Fig S2.** Relationship between the floral traits studied and the elevation, in meters above sea level. Both floral size (measured as flower weight) and the area of the standard petal increase with altitude (*P*= <0.001, *N*= 1124 flowers from 225 plants for flower size, and *P*= 0.04, *N*= 985 flowers from 225 plants for standard area). Colours in the figure indicate the different study sites.

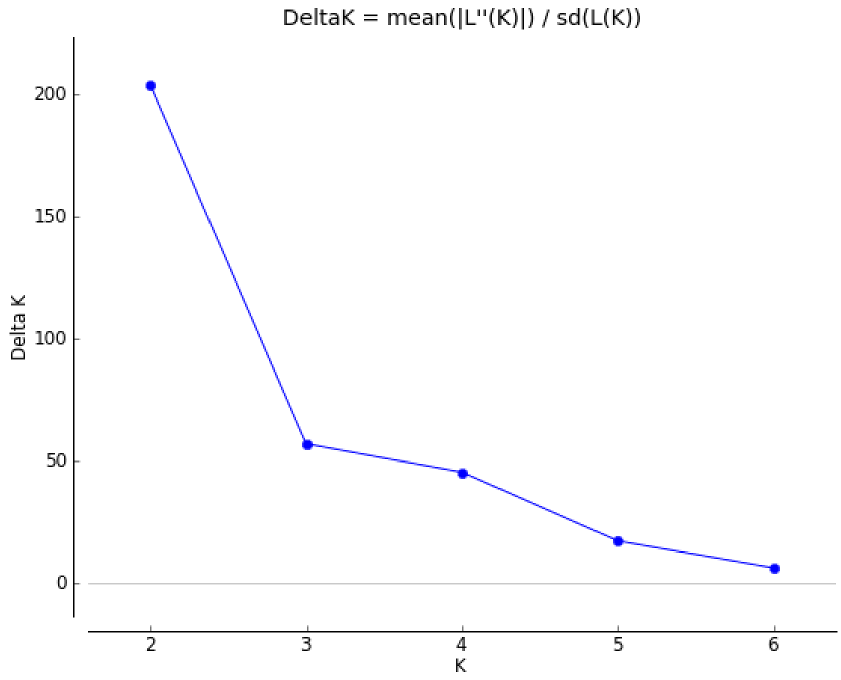


**Fig S3.** Results of population genetic structure and differentiation analysis for *Ulex parviflorus* localities. Evanno *et al.*'s (2005) method to identify the value of K that captures the highest level of structure suggests a maximum of K= 2 (i.e., the lowest limit of the method). That is, the six sites cannot be considered distinct populations.

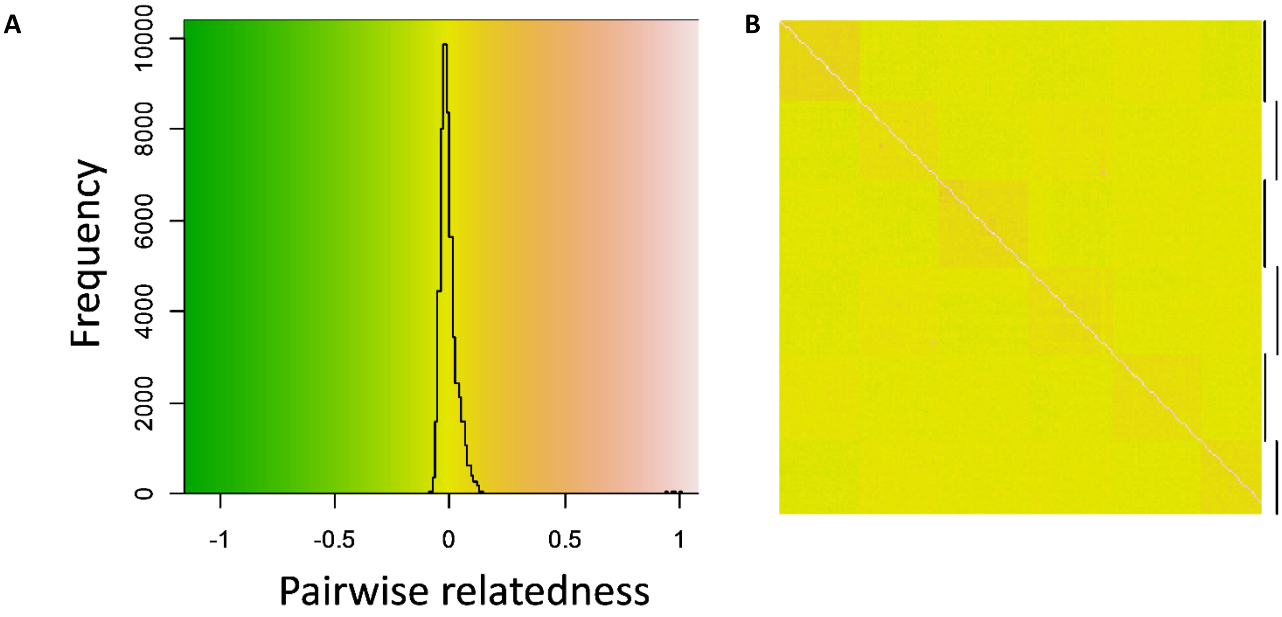


**Fig S4.** Estimates of pairwise genomic relatedness across all study localities. **A**) Histogram of relatedness values, and **B**) heatmap of estimates with colours corresponding to those in A. Black horizontal bars show the individuals from the six study localities.
